# Crowning the King of Fruits: Genomic Evidence of Population Structure and Relatedness in Malaysian Durians

**DOI:** 10.1101/2025.10.29.685476

**Authors:** Hung-Hui Chung, Li Yuan Liew, Adriana Batrisyia Mohd Faisal, Leonard Whye Kit Lim, Han Ming Gan

## Abstract

Durian (*Durio zibethinus*), often dubbed the King of Fruits, is a high-value tropical crop native to Southeast Asia and widely cultivated in Malaysia. Revered for its unique flavor yet polarizing for its pungent aroma, durian holds significant cultural and economic importance across the region. Despite this, genomic research on Malaysian durian cultivars remains limited. We performed whole-genome sequencing of several renowned and local durian cultivars, including Musang King (D197), Sultan (D24), Black Thorn (D200), Muar Gold (D101), and Red Prawn (D13) collected from distinct geographic regions across Malaysia. Illumina sequencing yielded approximately 20 to 30x coverage per sample, sufficient for single-nucleotide polymorphism (SNP) detection and population genomic analyses. Principal component analysis (PCA) based on genome-wide SNPs revealed visible population stratification and varying degrees of genetic relatedness among Malaysian durians. Subsequent KING kinship analysis indicated that most cultivars are relatively clonal, with the notable exception of the reference Musang King genome, which appeared genetically heterogeneous compared to other Musang King samples, a finding further supported by population structure inference from admixture analysis and ITS1-5.8S-ITS2 haplotyping. Notably, the Black Thorn cultivar showed no relatedness to Malaysian varieties but exhibited distant genetic similarity to the northern Thai Kan Yao variety. These results provide new genomic insights into the population structure, cultivar differentiation, and potential hybridization among Malaysian durians, forming a valuable foundation for future molecular marker development and breeding programs aimed at different variety authentication and genetic improvement.

## Introduction

Durian (*Durio zibethinus*) is a highly popular seasonal fruit in Malaysia, with strong domestic demand and growing export value, particularly to China (Figure 1). As a key contributor to Malaysia’s agricultural economy, durian holds both economic and cultural importance (Thorogood et al., 2022). Decades of selective breeding have produced numerous cultivars that differ in taste, texture, aroma, and bitterness, reflecting regional adaptation and consumer preference (Khoo, 2025). Despite Malaysia’s central role in durian domestication and cultivation, the first reference durian genome was published by Teh et al. (2017), using a specimen originally sourced from Pahang, Malaysia. The study offered insights into the molecular basis of durian’s distinctive aroma and established a well-annotated reference for future resequencing and comparative analyses.

**Figure 1.**
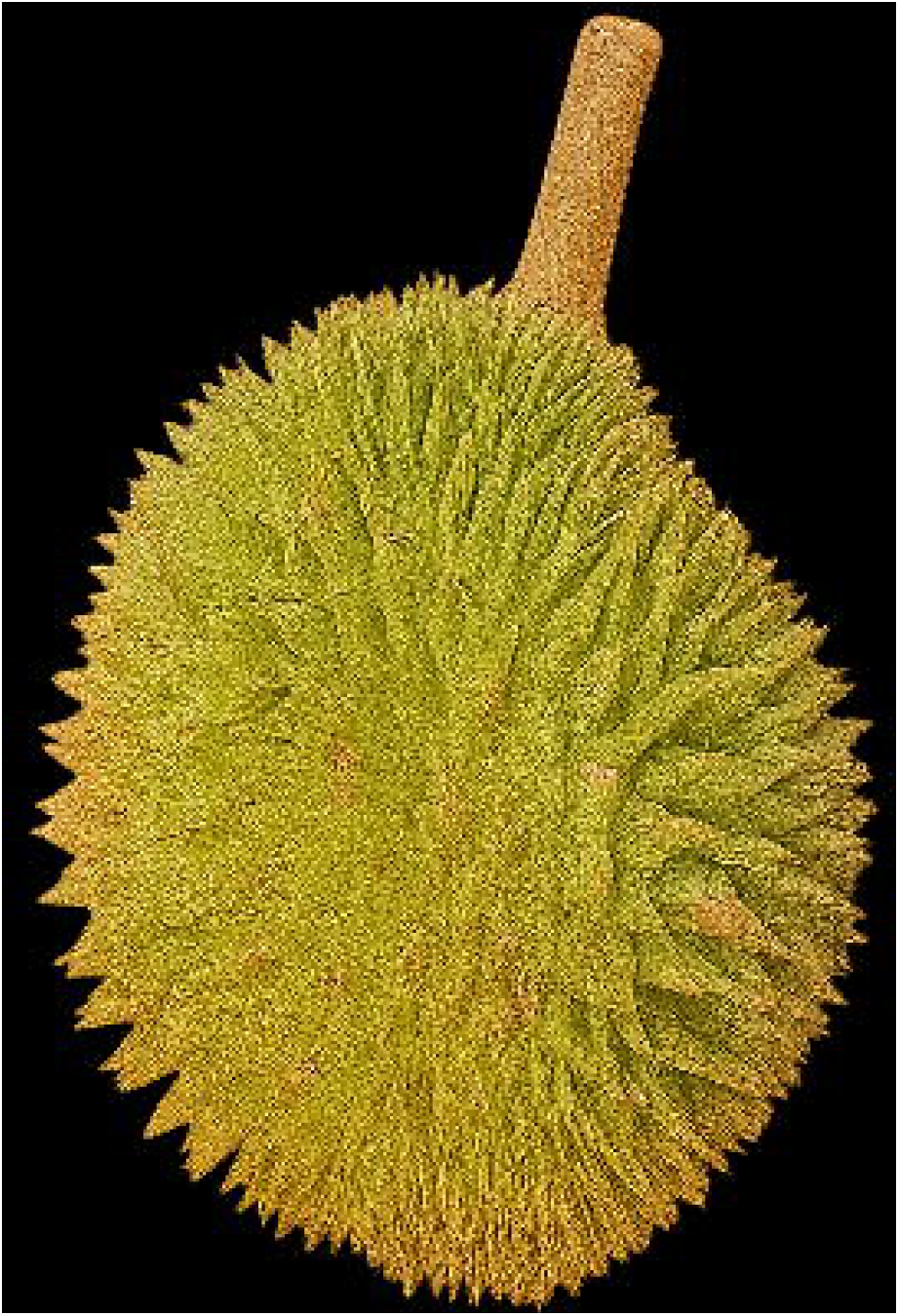
Durian fruit (kampung variety) from Sarawak, Malaysia, displayed for sale. The fruit shows the characteristic thorny husk and is fully ripe, ready to be opened for consumption.

Subsequent studies have expanded the genomic understanding of durian where Lin et al. (2022) performed ddRAD-seq analyses on durians cultivated in southern China, tracing their genetic origins to Southeast Asia. More recently, Zhong et al. (2025) performed whole-genome resequencing of over 100 durians planted in China and identified three genetic clusters, suggesting considerable genomic diversity among cultivated materials. However, because the accessions were all introduced and cultivated in China, with limited representation of native Southeast Asian germplasm, the study offers only a partial view of durian’s natural diversity. The genetic clustering observed may largely reflect recent propagation and planting histories rather than deep biological meaningful evolutionary differentiation. The weak correspondence between genetic clusters and geographic provenance, together with signs of reduced nucleotide diversity in one group, further suggests founder effects and a restricted genetic base among the Chinese-planted durians (Zhong et al., 2025)

On the contrary, in Thailand, three Thai durian genomes were sequenced, representing the Kradumthong, Monthong, and Puangmanee cultivars and similarly revealed substantial intraspecific variation among durian cultivars (Nawae et al., 2023). Building on this, another effort reported a chromosome-scale, haplotype-resolved assembly of the Thai cultivar Kan Yao, currently the most complete durian genome available (Ji et al., 2025).

Within Malaysia, however, population-level genomic research remains limited. The only relatively comprehensive durian genetic study to date used microsatellite markers to assess durian diversity across regions, revealing considerable genetic heterogeneity. Yet, the small number and modest informativeness of these loci restricted the resolution needed to distinguish among all cultivars (Siew et al., 2018). Beyond the Musang King reference genome, no other Malaysian durian genomes have been sequenced and/or reported, and none by a Malaysian-led research team. Given the deep cultural symbolism and economic significance of durian to Malaysia, this represents not merely a scientific gap but also a missed opportunity for national leadership in the genomic characterization of one of its most iconic crops.

In this study, we address this by generating whole-genome sequences for 14 durian samples representing major cultivars collected across Malaysia, including one from Sarawak, to capture the breadth of the country’s genetic and geographical diversity. Using Illumina whole-genome sequencing, we present the first population-scale genomic analysis of Malaysian durians, revealing patterns of genetic diversity and population stratification among key cultivars while reaffirming local genomic stewardship over this economically and culturally important species.

## Materials and Methods

### Sample collection and DNA Extraction for Whole Genome Sequencing

A total of 14 durian samples were collected across Malaysia for DNA extraction and whole-genome sequencing (Table 1). Approximately 150 mg of flesh or leaf tissue was homogenized and washed in sorbitol buffer (Inglis et al., 2018) to minimize polysaccharide and polyphenol carryover. Plant cell lysis was performed in a buffer containing 1% SDS, 25 mM Tris–HCl (pH 8.0), 10 mM EDTA, 15 mM NaCl and 2% PVP followed by incubation at 65 °C for 30 minutes. Subsequently, 0.5x volume of saturated potassium chloride was added to precipitate residual mucopolysaccharides (Sokolov, 2000). DNA was precipitated from the supernatant by adding 0.6x volume of isopropanol, centrifuged at maximum speed for 10 minutes, and the resulting DNA pellet was washed twice with 70% ethanol and resuspended in TE buffer.

**Table 1.** Summary of sequencing metrics, genome coverage and sampling information sorted by percentage of chloroplast-aligned reads.

| Sample ID | Total Mapped reads | Nuclear Genome Coverage (x) | Chloroplast Genome Coverage (x) | Chloroplast Reads (%) | Tissue Type | Preservation Method | Location | Reference |
| --- | --- | --- | --- | --- | --- | --- | --- | --- |
| KGSwk | 132369811 | 26 | 31579 | 22 | Flesh | Ethanol | Kuching, Sarawak | This study |
| MKSD | 176380995 | 35 | 36046 | 19 | Flesh | Frozen (-20C) | Sungai Dua, Pahang | This study |
| D101MT1 | 145065423 | 29 | 24548 | 16 | Flesh | Frozen (-20C) | Mantin, Negeri Sembilan | This study |
| D24SD | 204662301 | 41 | 34177 | 15 | Flesh | Frozen (-20C) | Sungai Dua, Pahang | This study |
| D101SD | 162594600 | 33 | 20460 | 11 | Flesh | Frozen (-20C) | Sungai Dua, Pahang | This study |
| KGSD | 186865730 | 37 | 16082 | 8 | Flesh | Frozen (-20C) | Sungai Dua, Pahang | This study |
| MKJ7 | 135854529 | 27 | 10374 | 7 | Leaf | Silica Gel | Johor | This study |
| MKRef | 127518986 | 26 | 6790 | 5 | Leaf | Unknown | Raub, Pahang | (Teh et al., 2017) |
| D101J2 | 154637389 | 31 | 5605 | 3 | Leaf | Silica Gel | Johor | This study |
| D13J3 | 184288643 | 37 | 5876 | 3 | Leaf | Silica Gel | Johor | This study |
| D13J4 | 147580309 | 30 | 4701 | 3 | Leaf | Silica Gel | Johor | This study |
| MKJ8 | 174925861 | 35 | 6117 | 3 | Leaf | Silica Gel | Johor | This study |
| BTJ6 | 178192152 | 36 | 4296 | 2 | Leaf | Silica Gel | Johor | This study |
| BTP | 163339770 | 33 | 4246 | 2 | Leaf | Silica Gel | Nibong Tebal, Pulau Pinang | This study |
| D101J1 | 162248528 | 32 | 3739 | 2 | Leaf | Silica Gel | Johor | This study |
| KanYao | 182566142 | 37 | 4454 | 2 | Leaf | Unknown | Thailand | (Ji et al., 2025) |

### Illumina Whole Genome Sequencing

Approximately 100 ng of high-quality genomic DNA was sheared to an average fragment size of 350 bp using a Bioruptor sonicator (Diagenode, Belgium).

Library preparation was performed using the NEBNext Ultra II DNA Library Preparation Kit (New England Biolabs, USA) following the manufacturer’s protocol. Briefly, fragmented DNA was end-repaired, followed by the addition of an adenine overhang and ligation of Illumina sequencing adapters. Indexed PCR amplification was then carried out to attach sample-specific barcodes and complete the Illumina adapter structure. The libraries were sequenced on an Illumina NovaSeq 6000 platform (Illumina, USA) using a paired-end 2 × 150 bp configuration, generating approximately 20 Gb of raw data per sample.

### Reference Mapping and Variant Calling

The reference durian nuclear genome of *Durio zibethinus* and the chloroplast genome from the *Durio zibethinus* Mon Thong variety (GenBank accession MT321069.1) (Shearman et al., 2020; Teh et al., 2017) were combined into a single reference assembly for read alignment. Raw Illumina reads were adapter- and quality-trimmed using fastp version 0.20.1 (Chen et al., 2018), retaining high-quality reads longer than 50 bp. Filtered reads were then aligned to the combined reference using BWA-MEM2 version 2.2.1 (Li, 2013). The resulting SAM files were sorted and converted to BAM format using SAMtools version 1.13 (Danecek et al., 2021). Variant calling was performed jointly on all aligned BAM files using bcftools v1.22 (Danecek et al., 2021). Variants were first generated with the mpileup command to produce uncompressed binary variant call output. The resulting variants were subsequently filtered using to retain only high-confidence (Q30), biallelic single-nucleotide polymorphisms (SNPs) with a minor allele frequency (MAF) ≥ 0.05 and missing data fraction ≤ 0.2. The final filtered VCF file was used for all downstream population genomic analyses, unless stated otherwise.

### Population Structure and Genetic Relatedness Analyses

Population structure among durian cultivars was first examined using principal component analysis (PCA) based on genome-wide single nucleotide polymorphisms (SNPs). High-quality SNP variants were converted from VCF to GDS format, and PCA was performed using the snpgdsPCA function from the SNPRelate v1.34.1 R package (Zheng et al., 2012). An admixture analysis was also conducted using ADMIXTURE v1.3 (Alexander and Lange, 2011), and the resulting ancestry proportions were visualized in a structure plot–style representation to reveal the genetic composition and possible admixture among cultivars. To further refine these relationships, kinship coefficients were subsequently computed using the KING algorithm implemented in PLINK v2.0 (Chang et al., 2015; Manichaikul et al., 2010). KING is designed to be robust to population stratification and allele frequency differences, providing a more accurate and unbiased estimation of true genetic relatedness.

### ITS1–5S–ITS2 Haplotype Reconstruction

Conventional variant calling approaches are unsuitable for the ITS1–5S–ITS2 region because, although the genome itself is diploid, this locus exists in multiple paralogous copies that vary within the same genome. As a result, standard diploid-based variant callers fail to capture the full haplotype diversity, and de novo assembly approaches often collapse these repetitive sequences into consensus contigs. To better resolve this complex region, a viral haplotyping–inspired approach was applied. First, a subsample of filtered pair-end reads (1 million reads) from sample BTP was assembled using MEGAHIT v1.2.9 (Li et al., 2015). Contigs containing the ITS region were identified using ITSx v1.1b (Bengtsson-Palme et al., 2013), which also extracted the full ITS1–5S–ITS2 region. Subsequently, 1 million filtered paired-end reads from each sample were then aligned to this reference sequence using BWA-MEM. Haplotype reconstruction was performed using CliqueSNV v2.0.3 (Knyazev et al., 2021), which infers local haplotypes and estimates their frequency based on patterns of co-occurring variants within the aligned reads. Finally, all reconstructed haplotypes from all samples were concatenated and re-clustered at 100% sequence identity and identical sequence length using CD-HIT v4.8.1 (Fu et al., 2012), producing a non-redundant haplotype set for downstream analyses.

## Results

### Durian flesh yields usable NGS data and higher chloroplast read counts

On average, 191 million (range: 166–222 million) raw paired-end reads were generated per sample (Supplemental Table 1), including those derived from durian flesh, which is typically more challenging due to the presence of polysaccharides and other PCR inhibitors. Interestingly, a higher proportion of reads from flesh samples aligned to the chloroplast genome compared to leaf samples, despite leaves being among the most chloroplast-rich tissues. Mapping analysis showed that 8–22% of reads from flesh samples aligned to the durian chloroplast genome, compared to only 2–7% from leaf samples (Table 1).

### PCA reveals population stratification among Malaysian durian cultivars

An initial set of 34.6 million SNPs was identified, of which 25.2 million high-quality SNPs remained following quality filtering. Principal component analysis (PCA) of genome-wide SNP data revealed notable genetic differentiation among the durian cultivars (Figure 2), depending on the principal component (PC) axes. Overall, the first four principal components captured the major dimensions of genetic variance and highlighted distinct population structure among the cultivars. PC1 effectively separated D101 (also known as IOI or Muar Gold) from all other cultivars, indicating strong genetic divergence and limited shared ancestry with the remaining groups. PC2 distinguished the premium Black Thorn (BT) and Musang King (MK) cultivars, with the first ever sequenced reference MK sample (MKRef) positioned slightly further along the axis. In contrast, PC3 primarily resolved D13 (Udang Merah), marking it as a distinct genetic lineage within the dataset. On the contrary, PC4 provided additional resolution by differentiating the Kampung durians (wild-type varieties). Interestingly, while both KGSD (Kampung from Selangor) and KGSwk (Kampung from Sarawak, Borneo) are grouped as “Kampung” types, their wide geographic separation is reflected genomically. PC4 clearly distinguishes the Sarawak sample from the Peninsular Malaysia one. However, the close clustering of D24 with KGSD is unexpected, given their distinct cultivar identities.

**Figure 2.**
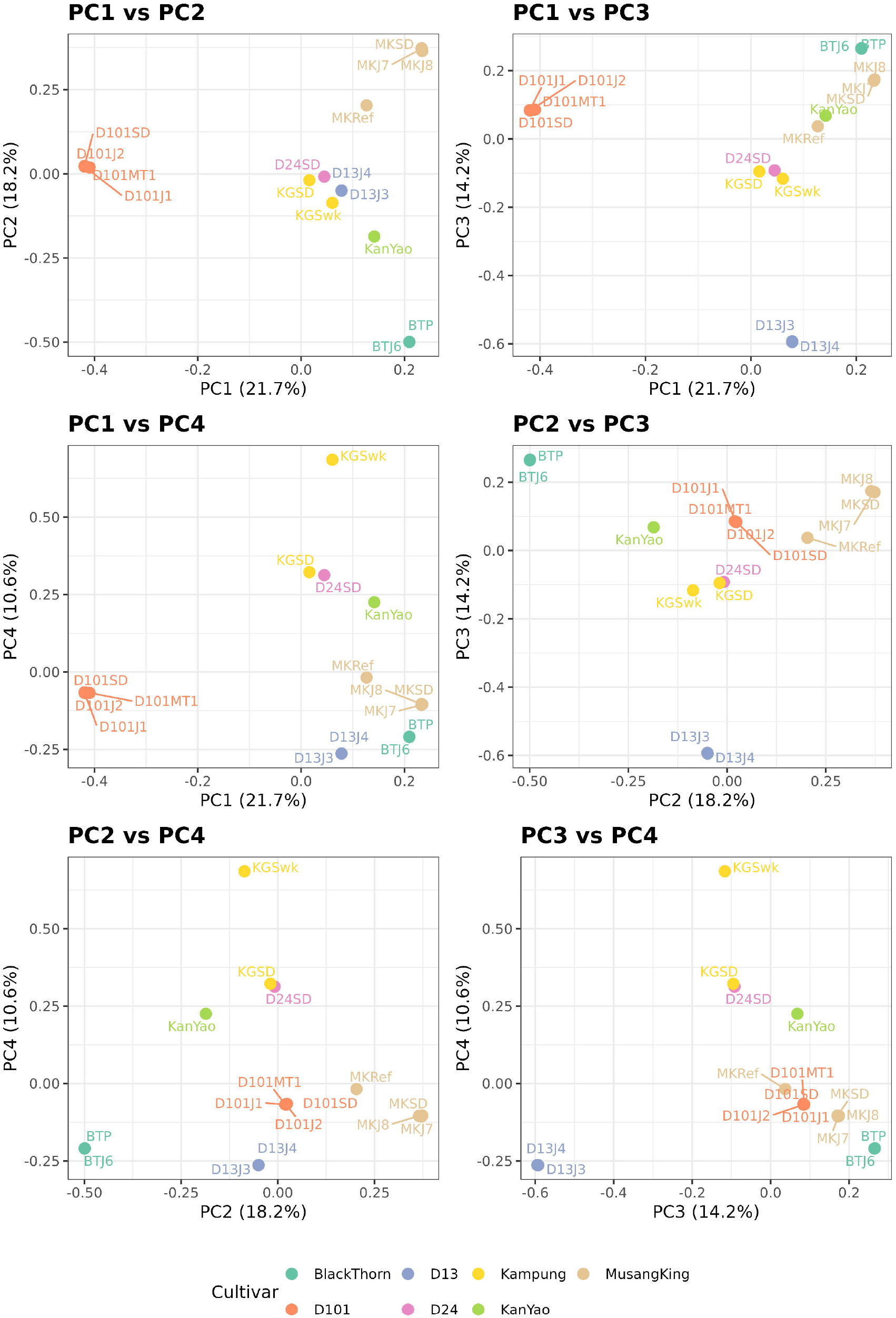
2D PCA plots showing different pairwise combinations of the first four principal components (PC1–PC4) derived from genome-wide SNP data. Each point represents an individual sample and is colored according to its durian cultivar. Clustering in each plot reflects variation along the specific PCs shown and does not necessarily represent absolute genetic relationships among samples. Percentage values on the axes indicate the proportion of total genetic variance explained by each principal component.

### The reference Musang King genome exhibits unexpected genetic heterogeneity compared to other Musang King samples

Admixture analysis based on genome-wide SNPs (Figure 3) revealed strong genetic differentiation and high cultivar purity among the major *Durio zibethinus* lineages. At K = 5, five distinct ancestry components were identified, corresponding to the principal cultivar groups: Musang King (MK), Black Thorn (BT), D24-Kampung, D13, and D101. Most samples showed nearly complete assignment to a single ancestry cluster, reflecting the high genetic homogeneity and clonal fidelity expected of graft-propagated cultivars. Notably, D24SD showed an ancestry composition identical to the Kampung cluster, indicating that the sampled tree is genetically indistinguishable from the Kampung variety at this resolution. All sequenced Musang King field samples (MKJ7, MKJ8, and MKSD) shared an almost identical genetic background despite being collected from different geographic regions, confirming the cultivar’s genomic consistency across locations. In contrast, the reference Musang King genome (MKRef) displayed a clear admixed profile, containing approximately 70–75% Musang King ancestry, with minor contributions from Black Thorn and D24-related components. Conversely, Black Thorn, D13, and D101 each formed clear, cultivar-specific groups with negligible admixture, underscoring their distinct breeding histories and minimal recent hybridization.

**Figure 3.**
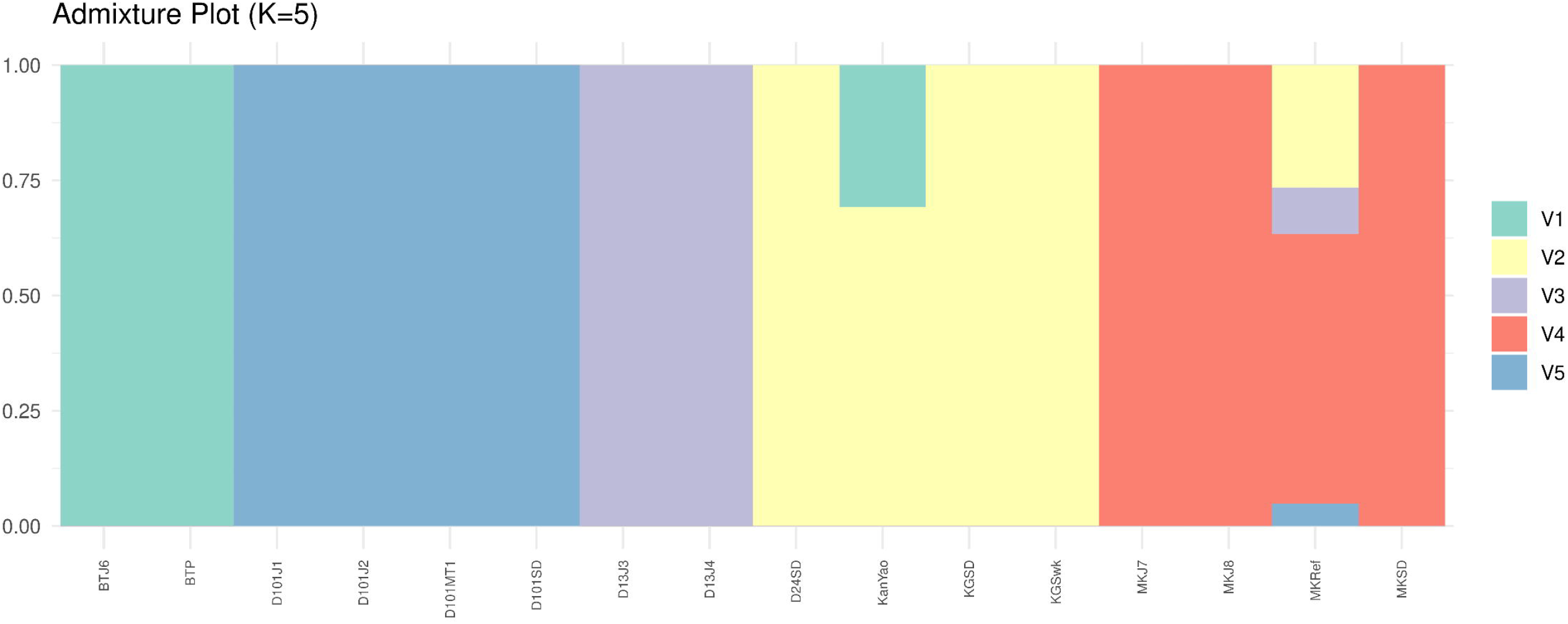
Structure plot based on Admixture analysis showing the proportional contribution of each ancestry component to individual samples. Analysis was performed with K = 5, which provides clear differentiation among the major durian cultivars. The height of each colored segment within a bar represents the estimated percentage of that ancestry in the corresponding sample.

### KING kinship supports clonal relationships for most cultivars and reveals unexpected distant links

Pairwise KING kinship coefficients (φ) were inferred from genome-wide SNP data (24.8–25.1 million SNPs) across 16zhang durian samples to robustly estimate recent co-ancestry while accounting for allele frequency variation. A kinship network was constructed with edges retained only for φ > 0.04, a moderately conservative threshold designed to capture both clonal and close pedigree relationships while excluding background relatedness. Edge thickness and color intensity were scaled by IBS0, the proportion of loci with zero alleles identical by state, such that lower IBS0 values denote greater genomic similarity (Figure 4). The resulting network revealed a central hub anchored by KGSD (a wild-type variety), connected to multiple samples. Musang King derivatives (MKJ7, MKJ8, MKSD, and MKRef) formed a tight clonal cluster characterized by low IBS0 values. Notably, the reference genome (MKRef) was connected to this cluster through slightly higher IBS0, suggesting close but non-clonal pedigree relationships. The D101 lineage (D101J1, D101J2, D101MT1, D101SD) formed a compact subclade radiating from KGSD, with uniformly low IBS0 (lightest edges), a pattern similarly observed for the D13 cluster. D24SD was linked only to D13 nodes and with relatively high IBS0, indicating distant but detectable co-ancestry. A separate Kan Yao/Black Thorn triad (KanYao, BTP, BTJ6) formed an independent module, with BTP–BTJ6 displaying very low IBS0 (clonal), while KanYao exhibited higher IBS0, indicating moderate divergence within this lineage. In contrast, KGSwk, originating from Sarawak, Malaysia, remained completely isolated from all other lineages.

**Figure 4.**
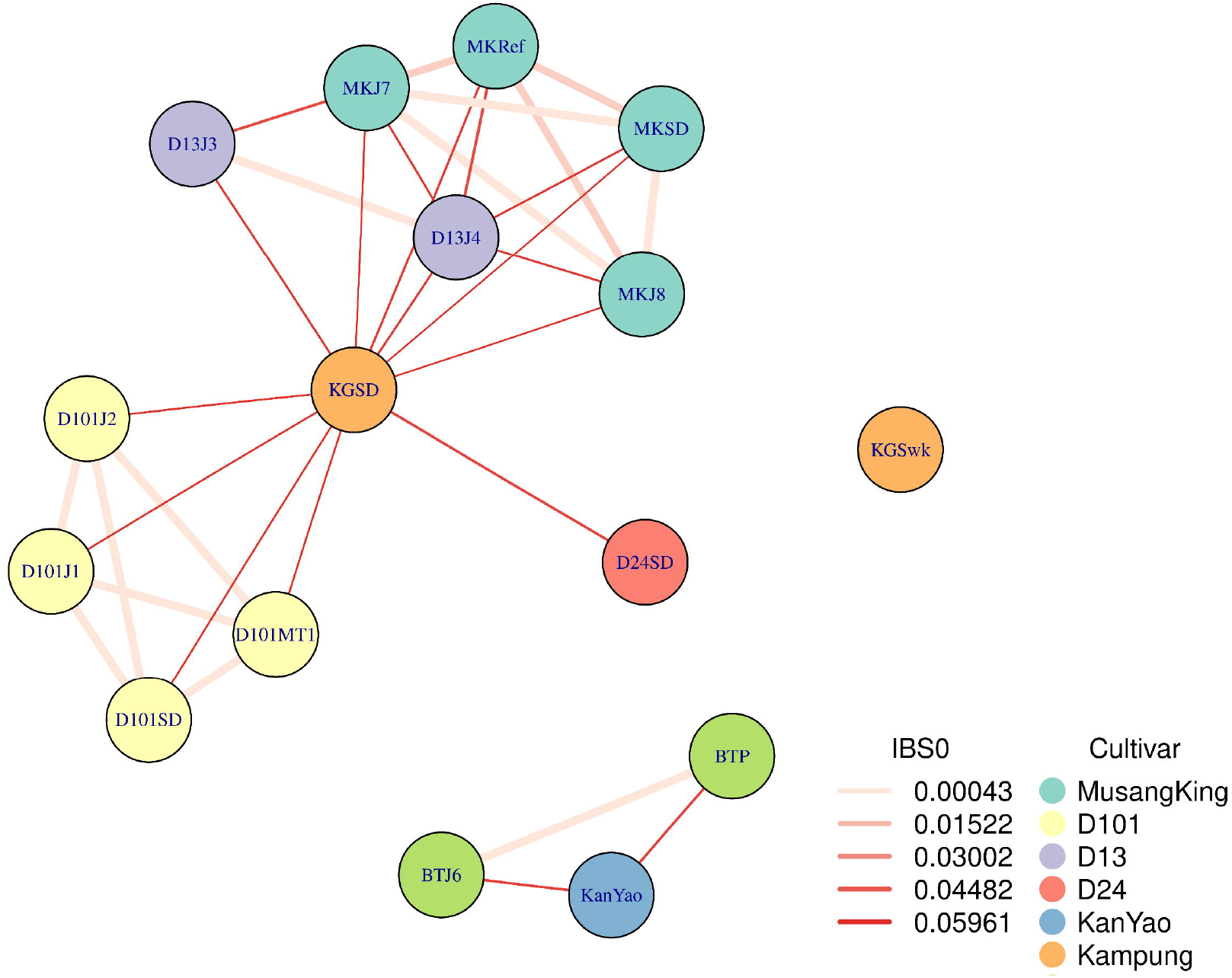
KING kinship network plot illustrating the pairwise kinship relationships among durian samples. Connections (edges) are drawn only between sample pairs with a KING kinship value ≥ 0.04. Edge colors represent the IBS0 values calculated from the KING analysis, where lower IBS0 indicates a higher proportion of shared alleles and higher IBS0 indicates fewer shared alleles. Nodes are colored according to cultivar.

### ITS1–5.8S–ITS2 haplotyping uncovers multiple paralogous variants

To further elucidate genetic relationships among cultivars and complement the high-resolution nuclear SNP analyses, we examined ITS1–5.8S–ITS2 haplotype composition derived from genome-skimming assemblies of the durian samples. Initial variant calling and haplotype reconstruction identified 26 haplotypes. After excluding those occurring at <5% within-sample frequency to minimize potential sequencing or assembly artifacts, 12 dominant haplotypes (Haplotype_0, _1, _2, _6, _7, _8, _9, _10, _11, _15, _21, and _22) were retained for downstream analyses. A presence/absence heatmap (Figure 5) revealed partial lineage-specific patterns. Haplotype_0, _2, and _6 were relatively common and shared among multiple cultivars, indicating limited diagnostic value. In contrast, Haplotype_1 was unique to the Black Thorn cluster (BTP and BTJ6), potentially serving as a strong diagnostic marker for this premium variety pending additional genomic sampling. Notably, the MKRef sample was distinguished by a private Haplotype_22 not detected in any other samples

**Figure 5.**
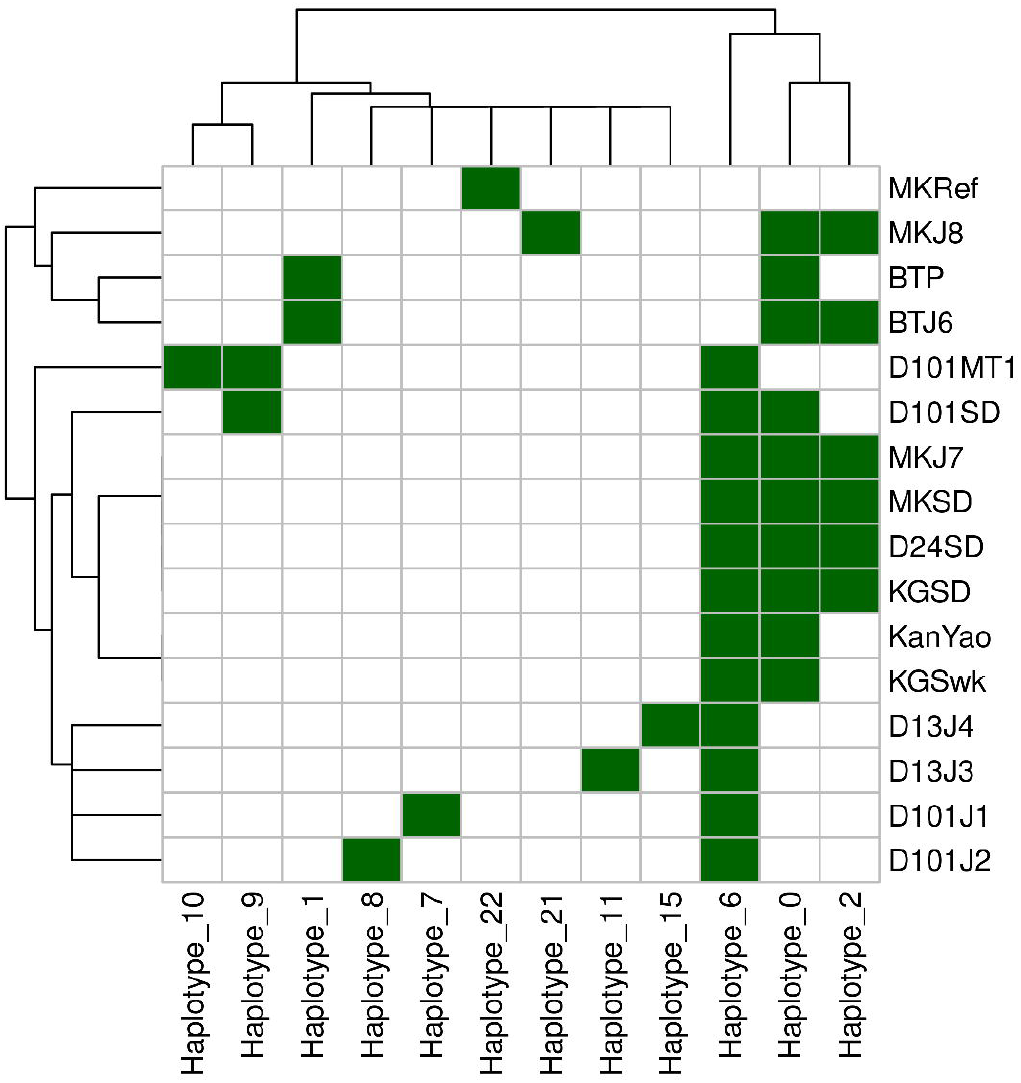
Presence/absence heatmap of ITS1–5.8S–ITS2 haplotypes across durian samples. Green indicates haplotype presence, and white indicates absence. Both samples and haplotypes were subjected to hierarchical clustering. A haplotype was considered present if its frequency exceeded 5% in a sample.

## Discussions

The successful generation of high-quality Illumina whole genome sequencing data from durian flesh, historically regarded as a challenging issue due to its high polysaccharide content (Ng et al., 2025), represents a promising advance for this species. This demonstrates that future genetic and genomic studies can be conducted directly from one of the most accessible parts of the fruit. Practically, this allows cultivar authenticity to be verified directly from the flesh sold at stalls or packaged for retail, offering a convenient and non-destructive alternative to leaf sampling that typically requires access to farms. Such an approach has strong implications for molecular authentication and traceability down to the level of individual fruit portions.

The stratification observed in genome-wide SNP-based principal component analysis indicates that Malaysian durian cultivars have been isolated long enough to develop cultivar-specific genomic signatures (Husin et al., 2018). These patterns reflect the unique breeding histories and regional selection pressures that shaped each variety. Interestingly, differentiation among certain lineages only became apparent in later principal components, suggesting that durian population structure cannot be fully represented by the first few components alone (Peloso and Lunetta, 2011; Zhong et al., 2025). Overreliance on these early components may therefore underestimate the true extent of cultivar stratification. One limitation of this study is the restricted number of samples, mainly due to the seasonal availability of fruits and limited access to rare or geographically localized cultivars.

The Musang King reference genome (MKRef), reported to have been collected from Raub in Pahang, exhibited distinct genetic characteristics compared with other Musang King accessions analyzed in this study. This divergence, consistently supported by principal component, ADMIXTURE, and ITS haplotype analyses, raises the possibility that MKRef represents a genetically mixed or non-pure sample. Given the popularity and economic importance of Musang King, this finding calls for a re-evaluation of the suitability of MKRef as the reference genome for this species. The Musang King samples included in our dataset were collected from several geographic regions and were genetically homogeneous relative to one another, which reinforces that the observed heterogeneity is unique to MKRef.

A key contribution of this work is the inclusion of two whole genome sequencing datasets for the Black Thorn cultivar, which currently commands a premium price higher than Musang King (Husin et al., 2018). Consistent with its historical origin in Penang (Siew et al., 2018), which is geographically close to Thailand, Black Thorn showed genetic affinity to the Thai cultivar Kan Yao rather than to other Peninsular Malaysian varieties. Future comparative analyses that include additional Thai cultivars may provide deeper insight into these regional relationships. It is worth noting that although three additional Thai durian genomes have recently been published, the raw sequencing data were not made available (Nawae et al., 2023) and therefore could not be incorporated into our population-level analyses. Even so, our results highlight the distinctiveness of Black Thorn, which also possesses a unique ITS1–5.8S–ITS2 haplotype profile that may serve as a useful molecular biomarker for cultivar identification.

The considerable variation observed among ITS haplotypes across cultivars highlights the dynamic evolution of rDNA loci in durian. Although ITS variation did not always align with genome-wide SNP patterns, this diversity is consistent with the known heterogeneity of multicopy rDNA regions (Wang et al., 2023; Xu et al., 2017). Because these regions are often collapsed during genome assembly and poorly represented in variant-based analyses, our approach, which applies a viral-inspired strategy (Knyazev et al., 2021) to reconstruct durian rDNA haplotypes, provides a new framework for exploring within-genome rDNA diversity. This method also demonstrates the potential for ITS-based markers to support durian species authentication, complementing established plant barcoding resources such as the ITS2 database (Banchi et al., 2020).

## Data Availability Statement

Raw paired-end FASTQ sequencing data for all durian samples generated in this study have been made publicly available under BioProject PRJNA1175461 Variant calling data, ITS1-5S-ITS2 haplotype sequences and KING kinship analysis output has been uploaded to Zenodo (https://zenodo.org/records/17479800)

## Acknowledgement

This work was supported by the AgriNXT 2023 Technology Challenge (Problem Statement AGRI 1: “Improving the accuracy and efficiency of durian variant identification”), funded by the Malaysian Communications and Multimedia Commission (MCMC). The authors also acknowledge the support of Bioeconomy Corporation and the Ministry of Agriculture and Food Security (KPKM) as organizing partners of the AgriNXT program. We extend our sincere thanks to Datuk Leow Cheok Kiang for providing the Black Thorn leaf samples and for facilitating access to his farm for sample collection.

## Notes

https://www.ncbi.nlm.nih.gov/bioproject/?term=PRJNA1175461

https://zenodo.org/records/17479800

### Competing Interest Statement

The authors have declared no competing interest.

## References

Alexander, D.H., Lange, K., 2011. Enhancements to the ADMIXTURE algorithm for individual ancestry estimation. BMC Bioinformatics 12, 246. 10.1186/1471-2105-12-246

Banchi, E., Ametrano, C.G., Greco, S., Stanković, D., Muggia, L., Pallavicini, A., 2020. PLANiTS: a curated sequence reference dataset for plant ITS DNA metabarcoding. Database 2020, baz155. 10.1093/database/baz155

Bengtsson-Palme, J., Ryberg, M., Hartmann, M., Branco, S., Wang, Z., Godhe, A., De Wit, P., Sánchez-García, M., Ebersberger, I., de Sousa, F., Amend, A., Jumpponen, A., Unterseher, M., Kristiansson, E., Abarenkov, K., Bertrand, Y.J.K., Sanli, K., Eriksson, K.M., Vik, U., Veldre, V., Nilsson, R.H., 2013. Improved software detection and extraction of ITS1 and ITS2 from ribosomal ITS sequences of fungi and other eukaryotes for analysis of environmental sequencing data. Methods Ecol. Evol. 4, 914–919. 10.1111/2041-210X.12073

Chang, C.C., Chow, C.C., Tellier, L.C., Vattikuti, S., Purcell, S.M., Lee, J.J., 2015. Second-generation PLINK: rising to the challenge of larger and richer datasets. GigaScience 4, 7. 10.1186/s13742-015-0047-8

Chen, S., Zhou, Y., Chen, Y., Gu, J., 2018. fastp: an ultra-fast all-in-one FASTQ preprocessor. Bioinformatics 34, i884–i890. 10.1093/bioinformatics/bty560

Danecek, P., Bonfield, J.K., Liddle, J., Marshall, J., Ohan, V., Pollard, M.O., Whitwham, A., Keane, T., McCarthy, S.A., Davies, R.M., Li, H., 2021. Twelve years of SAMtools and BCFtools. GigaScience 10, giab008. 10.1093/gigascience/giab008

Fu, L., Niu, B., Zhu, Z., Wu, S., Li, W., 2012. CD-HIT: accelerated for clustering the next-generation sequencing data. Bioinformatics 28, 3150–3152. 10.1093/bioinformatics/bts565

Husin, N.A., Rahman, S., Karunakaran, R., Bhore, S.J., 2018. A review on the nutritional, medicinal, molecular and genome attributes of Durian (Durio zibethinus L.), the King of fruits in Malaysia. Bioinformation 14, 265–270. 10.6026/97320630014265

Inglis, P.W., Pappas, M. de C.R., Resende, L.V., Grattapaglia, D., 2018. Fast and inexpensive protocols for consistent extraction of high quality DNA and RNA from challenging plant and fungal samples for high-throughput SNP genotyping and sequencing applications. PLoS ONE 13, e0206085. 10.1371/journal.pone.0206085

Ji, X., Zhong, Y., Zheng, D., Xie, S., Shi, M., Wang, X., Liu, F., Feng, X., Wang, H., 2025. Chromosome-scale haploid genome assembly of Durio zibethinus KanYao. Sci. Data 12, 384. 10.1038/s41597-025-04656-y

Khoo, G.C., 2025. Durian matters. Continuum 39, 211–217. 10.1080/10304312.2024.2421883

Knyazev, S., Tsyvina, V., Shankar, A., Melnyk, A., Artyomenko, A., Malygina, T., Porozov, Y.B., Campbell, E.M., Switzer, W.M., Skums, P., Mangul, S., Zelikovsky, A., 2021. Accurate assembly of minority viral haplotypes from next-generation sequencing through efficient noise reduction. Nucleic Acids Res. 49, e102. 10.1093/nar/gkab576

Li, D., Liu, C.-M., Luo, R., Sadakane, K., Lam, T.-W., 2015. MEGAHIT: an ultra-fast single-node solution for large and complex metagenomics assembly via succinct de Bruijn graph. Bioinformatics 31, 1674–1676. 10.1093/bioinformatics/btv033

Li, H., 2013. Aligning sequence reads, clone sequences and assembly contigs with BWA-MEM. ArXiv13033997 Q-Bio.

Manichaikul, A., Mychaleckyj, J.C., Rich, S.S., Daly, K., Sale, M., Chen, W.-M., 2010. Robust relationship inference in genome-wide association studies. Bioinformatics 26, 2867–2873. 10.1093/bioinformatics/btq559

Nawae, W., Naktang, C., Charoensri, S., U-thoomporn, S., Narong, N., Chusri, O., Tangphatsornruang, S., Pootakham, W., 2023. Resequencing of durian genomes reveals large genetic variations among different cultivars. Front. Plant Sci. 14. 10.3389/fpls.2023.1137077

Ng, W.L., Wong, X.J., Ramaiya, S.D., Sabran, S., Hung, B.M., Lee, S.Y., 2025. Extraction of Useful DNA from Different Parts of the Durian (Durio zibethinus) Fruit. Pertanika J. Trop. Agric. Sci. 48. 10.47836/pjtas.48.4.03

Peloso, G.M., Lunetta, K.L., 2011. Choice of population structure informative principal components for adjustment in a case-control study. BMC Genet. 12, 64. 10.1186/1471-2156-12-64

Shearman, J.R., Sonthirod, C., Naktang, C., Sangsrakru, D., Yoocha, T., Chatbanyong, R., Vorakuldumrongchai, S., Chusri, O., Tangphatsornruang, S., Pootakham, W., 2020. Assembly of the durian chloroplast genome using long PacBio reads. Sci. Rep. 10, 15980. 10.1038/s41598-020-73549-4

Siew, G.Y., Ng, W.L., Tan, S.W., Alitheen, N.B., Tan, S.G., Yeap, S.K., 2018. Genetic variation and DNA fingerprinting of durian types in Malaysia using simple sequence repeat (SSR) markers. PeerJ 6, e4266. 10.7717/peerj.4266

Sokolov, E.P., 2000. An improved method for DNA isolation from mucopolysaccharide-rich molluscan tissues. J. Molluscan Stud. 66, 573–575. 10.1093/mollus/66.4.573

Teh, B.T., Lim, K., Yong, C.H., Ng, C.C.Y., Rao, S.R., Rajasegaran, V., Lim, W.K., Ong, C.K., Chan, K., Cheng, V.K.Y., Soh, P.S., Swarup, S., Rozen, S.G., Nagarajan, N., Tan, P., 2017. The draft genome of tropical fruit durian (Durio zibethinus). Nat. Genet. 49, 1633–1641. 10.1038/ng.3972

Thorogood, C.J., Ghazalli, M.N., Siti-Munirah, M.Y., Nikong, D., Kusuma, Y.W.C., Sudarmono, S., Witono, J.R., 2022. The king of fruits. PLANTS PEOPLE PLANET 4, 538–547. 10.1002/ppp3.10288

Wang, W., Zhang, X., Garcia, S., Leitch, A.R., Kovařík, A., 2023. Intragenomic rDNA variation - the product of concerted evolution, mutation, or something in between? Heredity 131, 179–188. 10.1038/s41437-023-00634-5

Xu, B., Zeng, X.-M., Gao, X.-F., Jin, D.-P., Zhang, L.-B., 2017. ITS non-concerted evolution and rampant hybridization in the legume genus Lespedeza (Fabaceae). Sci. Rep. 7, 40057. 10.1038/srep40057

Zheng, X., Levine, D., Shen, J., Gogarten, S.M., Laurie, C., Weir, B.S., 2012. A high-performance computing toolset for relatedness and principal component analysis of SNP data. Bioinformatics 28, 3326–3328. 10.1093/bioinformatics/bts606

Zhong, Y., Feng, L., Deng, H., Ji, X., Zhang, J., Sun, Y., Lin, P., Qiao, Y., Xie, S., Wang, H., Guo, L., Feng, X., 2025. Genomic resequencing reveals genetic diversity, population structure, and core collection of durian germplasm. Commun. Biol. 8, 1273. 10.1038/s42003-025-08715-3

